# Longitudinal standards for mid-life cognitive performance: Identifying abnormal within-person changes in the Wisconsin Registry for Alzheimer’s Prevention

**DOI:** 10.1101/229146

**Authors:** Rebecca L. Koscik, Erin M. Jonaitis, Lindsay R. Clark, Kimberly D. Mueller, Samantha L. Allison, Carey E. Gleason, Richard Chappell, Bruce P. Hermann, Sterling C. Johnson

## Abstract

**Objective:** A major challenge in cognitive aging is differentiating preclinical disease-related cognitive decline from changes associated with normal aging. Neuropsychological test authors typically publish single time-point norms, referred to here as *unconditional* standards or reference values. However, detecting significant change requires longitudinal, or *conditional* XSreference values, created by modeling cognition as a function of prior performance. Our objectives were to create, depict, and examine preliminary validity of unconditional and conditional reference values for ages 40-75 on neuropsychological tests of memory and executive function.

**Method:** We used quantile regression to create growth-curve-like models of performance on tests of memory and executive function using participants from the Wisconsin Registry for Alzheimer’s Prevention. Unconditional and conditional models accounted for age, sex, education, and verbal ability/literacy; conditional models also included past performance on and number of prior exposures to the test. Models were then used to estimate individuals’ unconditional and conditional percentile ranks for each test. We then examined how low performance on each test (operationalized as <7th percentile) related to consensus-conference-determined cognitive statuses, and subjective impairment.

**Results:** Participants with low performance according to the reference values were more likely to receive an abnormal cognitive diagnosis at the current visit (but not later visits). Low performance was also linked to subjective and informant reports of worsening memory function.

**Conclusions:** Methods are needed to identify significant within-person cognitive change. The unconditional and conditional reference-development methods described here have many potential uses in research and clinical settings.

## Introduction

A major challenge in the field of cognitive aging is differentiating disease-related cognitive change from the more gradual decline expected in normal aging, which is particularly difficult during the preclinical stage of cognitive decline. Forthcoming diagnostic guidelines for Alzheimer’s disease (AD) posit such a period – --Clinical Stage 2”-- which encompasses significant within person change from a previous level of functioning that is not yet severe enough to be categorized as mild cognitive impairment (MCI; Jack et al., 2017). However, precise guidance on how this preclinical change should be operationalized is not yet established.

While most normed neuropsychological instruments intended for use in geriatric populations publish performance norms for several age bands (e.g., Steinberg, Bieliauskas, Smith, Ivnik, & Malec, 2005), with or without adjustments for relevant demographic features such as sex, education, or intelligence, they rarely account for a patient’s prior cognitive performance on the instrument itself. The clinician or researcher must resort to using reliable change or deviation estimates that may not fully account for underlying age-associated change or practice effects. Norms that adjust for a test-taker’s prior performance as well as demographic features would improve the validity of the interpretation in such circumstances. One approach to this problem has been the use of a regression-based approach to predicting change (Attix et al., 2009; Crawford, Garthwaite, Denham, & Chelune, 2012; Duff et al., 2005; B. P. Hermann et al., 1996; Maassen, Bossema, & Brand, 2009). In this approach, participants’ demographics and baseline scores are used to calculate expected scores on follow-up tests. The difference between predicted and observed values is then compared against an estimate of the standard error of prediction (*SE*_*p*_); individual scores exceeding some threshold (e.g. ±1.5*SE*_*p*_) are considered evidence of reliable change. Although influential, this approach assumes that the relationships between predictors and test scores are constant across all predictor strata. It is also rather indirect, since it relies on estimating the mean to understand individual performance that is far from the mean.

Methodologies first developed for anthropometric indices (e.g., height and weight) provide an alternative approach. The first reference curves for height were published over a century ago using heights of Massachusetts schoolchildren (Bowditch, 1891; Cole, 2012). Similar curves are used still today to evaluate children’s development (WHO Multicentre Growth Reference Study Group, 2006). Although reference curves are commonly termed growth curves, in reality they are often produced using cross-sectional anthropometric data, and provide little information about individual height trajectories (Tanner, Whitehouse, & Takaishi, 1966). When considering the growth of an individual child, the quantity of most interest is typically how usual or unusual their stature is now, given previous measurements. We refer to reference curves developed in this way as *conditional*, in contrast with the *unconditional* reference curves produced with cross-sectional information only. Conditional curves can be developed by regressing stature at time *t*_*j*_ on stature at time *t*_*j*-1_, controlling for the interval between observations (Berkey, Reed, & Valadian, 1983; Cameron, 1980; Cole, 1995; Healy, 1974). The earliest analyses of conditional curves used parametric methods, making them conceptually similar to the regression-based methods described by Maassen (2009); however, more recent work has instead used quantile regression, which requires fewer distributional assumptions (Wei, Pere, Koenker, & He, 2006) and gives directly relevant results which standard regression models of means do not. Through these methods, researchers can obtain curves describing the expected median cognitive trajectory, as well as trajectories for more extreme quantiles that denote unexpected loss or gain. Selection of quantiles is based upon their intended use and the amount of information available, as larger datasets enable the estimation of more extreme quantiles than smaller ones.

Conditional reference curves have recently been applied to the field of cognitive aging. Cheung and colleagues demonstrated the development of unconditional and conditional references using the Mini Mental State Exam (MMSE; Cheung et al., 2015). Using quantile regression, they established smooth, age-linked reference curves for several percentiles of interest adjusting only for baseline covariates. A second set of per-person conditional reference curves was then created, adjusting both for covariates and previous MMSE performance. The conditional references provided a much narrower scope for normative performance than the unconditional references, as well as a clear visual aid for identifying potentially concerning change.

In this work we extend the conditional reference methodology of Cheung and colleagues to the Wisconsin Registry for Alzheimer’s Prevention (WRAP) dataset. WRAP is a longitudinal cohort study of middle-aged and older adults who complete cognitive testing at regular intervals. This cohort is enriched with risk for Alzheimer’s disease (AD) due to parental family history (S. C. Johnson et al., n.d.; Sager, Hermann, & La Rue, 2005). Their mean age at first visit was 54, making WRAP an ideal population in which to examine preclinical cognitive decline. Our goals in this paper were to: (1) extend the unconditional and conditional reference methods using a range of tests known to be sensitive to preclinical decline; (2) develop a graphical tool for contextualizing individual performance over time; (3) begin reviewing validity evidence for the method by examining how abnormal conditional performance (ACP) relates to cognitive status and subjective functioning; and last, (4) explore differences between individuals that are flagged for abnormal unconditional and conditional performance. The overarching goal of these analyses is to develop a procedure that can be used in this and other cohorts to identify Stage 2 (preclinical) decline.

## Methods

### Participants

The WRAP cohort currently includes neuropsychological data from 1561 participants who enrolled at midlife (~40-65 years of age) and were free of dementia at baseline. Follow-up visits are conducted at two-to four-year intervals. Participant retention is approximately 81%; median follow-up is 9 years for active participants (S. C. Johnson et al., n.d.). This ongoing study is conducted in compliance with ethical principles for human subjects research defined in the Declaration of Helsinki, including review and approval by the University of Wisconsin Institutional Review Board, and the provision of informed consent by all participants.

#### Inclusion/exclusion criteria

When constructing growth curves, whether conditional or unconditional, the question of whom to include in the sample is paramount (Cole, 2012; Corvalan, 2014). A *reference curve* aims to describe typical growth, whereas a *standard* uses a sample selected for optimal health (*e.g.*, WHO Multicentre Growth Reference Study Group, 2006; J. Xu, Luntamo, Kulmala, Ashorn, & Cheung, 2014). The protracted preclinical phase and variable age of onset associated with AD-related dementia and other dementias make it difficult to apply these strategies for sample selection, because it is not clear which members of a given sample are truly free of disease. Instead, many studies exclude only people already evincing clinically-significant impairment, such as those having a diagnosis of probable AD dementia or Parkinson’s disease (Kenny et al., 2013) and those with baseline scores on other neuropsychological tests suggestive of MCI (Cheung et al., 2015). Because our exclusion criteria included a clinical/neurocognitive component, we refer to the curves we describe in this paper as standards.

For these analyses, we selected the subset of WRAP participants who were free of clinical MCI or dementia through their first two study visits and were free of neurological conditions that could affect cognition. Exclusionary criteria (n) included: consensus diagnosis of MCI or dementia (n=14), or self-reported diagnosis of epilepsy, stroke, multiple sclerosis, or Parkinson’s disease (n=58), before Visit 3; had not yet completed Visit 3 (n=376); had missing outcome or predictor data (n=10); or were outside target age range (40-75) for any of the first three visits (n=14). After exclusions, 1089 participants were included in the standards development sample.

### Measures

#### Cognitive and clinical outcomes

At each visit, participants complete a comprehensive neuropsychological battery (details in S. C. Johnson et al., n.d.). For these analyses, we created standards for the following tests and items: Rey Auditory Verbal Learning Test (AVLT; Schmidt, 1996), learning trials sum and delayed recall trial; Trail-Making Test (TMT; Reitan, 1958), part A and part B; WAIS-III Digit Span (Wechsler, 1997), forward and backward; Stroop Test (Trenerry, 1989), Color and Color-Word trials. These tests were selected based on the sensitivity of these domains to early cognitive impairment (Hedden, Oh, Younger, & Patel, 2013) and the completeness of data, as all were administered at the participant’s baseline. We also created standards for discrepancy scores calculated as follows: AVLT, delayed recall minus the last learning trial (trial 5); Trail-Making Test, part A minus part B; Digit Span, backward minus forward; and Stroop, color-word minus color. These discrepancy scores were of interest to us based on earlier evidence from our sample that intraindividual cognitive variability may be indicative of cognitive deterioration (Koscik et al., 2016).

Informant-based assessments of clinical symptoms were also collected, including the Quick Dementia Rating System (QDRS; Galvin, 2015) and/or the Clinical Dementia Rating Scale (CDR; Morris, 1997), combined as described in Berman (Berman et al., 2017; range = 0 to 3, 0.5 indicates MCI), and the Informant Questionnaire on Cognitive Decline in the Elderly, or IQCODE (Jorm & Jacomb, 1989; range = 16 to 80, 48 represents no change, higher scores indicate worsening functioning). Subjective complaint measures included two items representing participants’ self-report of memory functioning: “Do you think you have a problem with your memory?” (0=no, 1=yes; Don’t know coded to missing); and “Overall, how would you rate your memory in terms of the kinds of problems that you have?” (Likert scale range from 1=“Major problems” to 7=“No Problems”) (Memory Functioning Questionnaire; Gilewski, Zelinski, & Schaie, 1990).

#### Cognitive status determination

For research purposes, participant cognitive status was determined after each study visit via a consensus conference review of cognitive performance, medical history, and other factors (for details, see S. C. Johnson et al., n.d.; Koscik et al., 2016). Cognitive statuses included: Cognitively Normal; Early MCI (i.e., scores that are low compared to our internal robust norms but which do not cross objective thresholds for MCI); MCI; Dementia; or Impaired Not MCI. This latter category is assigned to a small number of participants whose history suggests longstanding impairment (e.g., history of learning disability) rather than a more recently acquired impairment. We included these participants in the standards development sample since they represent normal variation in the population.

### Statistical methods

#### Software

Data management tasks were done using R (R Core Team, 2017) and SAS software version 9.4. Analyses were conducted in R, and documented using RStudio (RStudio Team, 2016) and knitr (Xie, 2017).

#### Standards development

We created two sets of standards: *unconditional standards*, which summarize the distribution of each outcome within demographic strata, but do not consider previous measurements of that outcome; and *conditional standards*, which take into account both demographics and past performance on that test. To build both sets, we constructed regression quantiles using the R package quantreg (Koenker, 2017; R Core Team, 2017). We selected nine quantiles of interest to include the median, the 25th and 75th percentiles, and three quantiles in each tail which, for a normally-distributed outcome, correspond to approximately ±1, ±1.5, and ±2 standard deviations away from the mean (2%, 7%, 16%, 25%, 50%, 75%, 84%, 93%, 98%). For each outcome, preliminary unconditional models including only linear and quadratic age terms were constructed for the selected quantiles. If the quadratic term was nominally significant (p<.05) for at least two quantiles, it was retained in the model for all percentiles.

Following model selection, we constructed unconditional standards for each cognitive outcome using selected age terms plus three categorical covariates: sex (0=male, 1=female), college completion (0=no bachelor’s degree, 1=has degree), and baseline WRAT reading score (Wilkinson, 1993; here categorized as 0=66-89, 1=90-99, 2=100-109, 3=110-120), included as a proxy for education quality and verbal ability (Manly, Touradji, Tang, & Stern, 2003). We then modeled conditional standards by further controlling for an individual’s mean score on a given outcome at Visits 1 and 2, along with their number of prior test exposures. For both conditional and unconditional standards, we constructed regression quantiles for all percentiles from 1 to 99. Estimated subject-specific conditional and unconditional percentiles were then derived by comparing each true score to the 99 predicted regression quantiles and selecting the quantile with the minimum absolute error. Cluster-robust bootstrap standard errors were used to control for the inclusion of multiple measurements per subject (Hagemann, 2017).

#### Abnormal unconditional and conditional performance

For any given visit and cognitive test, a participant’s performance was referred to as abnormal unconditional performance (AUP) if the test score fell below the 7th unconditional percentile. For a normally-distributed outcome, this percentile corresponds to approximately -1.5 SD below the expected mean for that stratum, a cutoff we and others have used in previous work evaluating MCI (e.g., Clark et al., 2016; Cook & Marsiske, 2006). Similarly, from Visit 3 onward, participants whose score fell below the 7th *conditional* percentile were flagged as exhibiting abnormal conditional performance (ACP) at that visit. Henceforth, each test score was associated with two binary variables indicating its abnormality compared to unconditional (0=normal, 1=AUP) or conditional (0=normal, 1=ACP) standards. A participant flagged for ACP or AUP at one visit for a given test was not necessarily flagged on the same test at the next visit.

#### Graphical tool

We developed a graphical tool to contextualize individual performance over time. Built on ggplot2 (Wickham, 2009), this module plots a participant’s individual test scores over time against a series of age-based curves representing unconditional regression quantiles for that participant’s sex, education, and literacy level. Test scores receiving ACP flags are circled. We present three cases in the results to illustrate use of the graphical tool.

#### Construct validity

To assess whether ACP on a given test measured increased risk of abnormal cognitive status (one aspect of construct validity), we asked two questions: Do our quantile-regression-based abnormal conditional performance (ACP) indicators at a given visit correlate with cognitive status at the *same visit*? And does ACP at an early visit, controlling for AUP, predict a person’s cognitive status at the *last visit*? To answer these questions, we constructed generalized linear models for each outcome, using data from Visit 3 onward (the first visit for which ACP information was available). For models linking ACP to concurrent diagnoses, we estimated the relationship using generalized estimating equations for ordinal outcomes (`multgee` package; Touloumis, 2015). For the question relating first-available ACP and AUP indicators to future diagnoses, we estimated the relationship using ordinal regression (`MASS` package, function `polr`; Venables & Ripley, 2002). For these analyses, we eliminated anyone whose cognitive status was Impaired Not MCI.

Evidence regarding the utility of subjective complaints in identifying cognitive impairment is mixed (Roberts, Clare, & Woods, 2009). However, people’s complaints may indicate that they have noticed a drop in their own performance that still leaves them above conventional thresholds for impairment, in which case we would expect ACP to be associated with these subjective measures. To test this, we modeled ACP as a function of each of three subjective measures using generalized estimating equations for binary outcomes (`geepack` package; Højsgaard, Halekoh, & Yan, 2006).

Each of the above questions regarding cognitive status and subjective cognitive complaints involved constructing twelve models (one for each of eight test scores, plus one for each of four discrepancy scores); for each set of twelve, we adjusted p-values for multiple comparisons using the Benjamini-Hochberg method for controlling the false discovery rate (Benjamini & Hochberg, 1995). This method assumes that outcomes are either independent or positively dependent, an assumption these analyses generally met (minimum pairwise *r* = −.07).

#### Joint distribution of ACP and AUP indicators

For all outcomes, we categorize participants’ Visit 3 performance as follows: normal by both standards; AUP only; ACP only; and both ACP and AUP. We expected that approximately 6-7% should meet criteria for each, but had no a priori hypothesis about the overlap between the two.

## Results

### Participant characteristics

Baseline characteristics for this sample are shown in Table 1 overall and by sex.

**Table 1.**
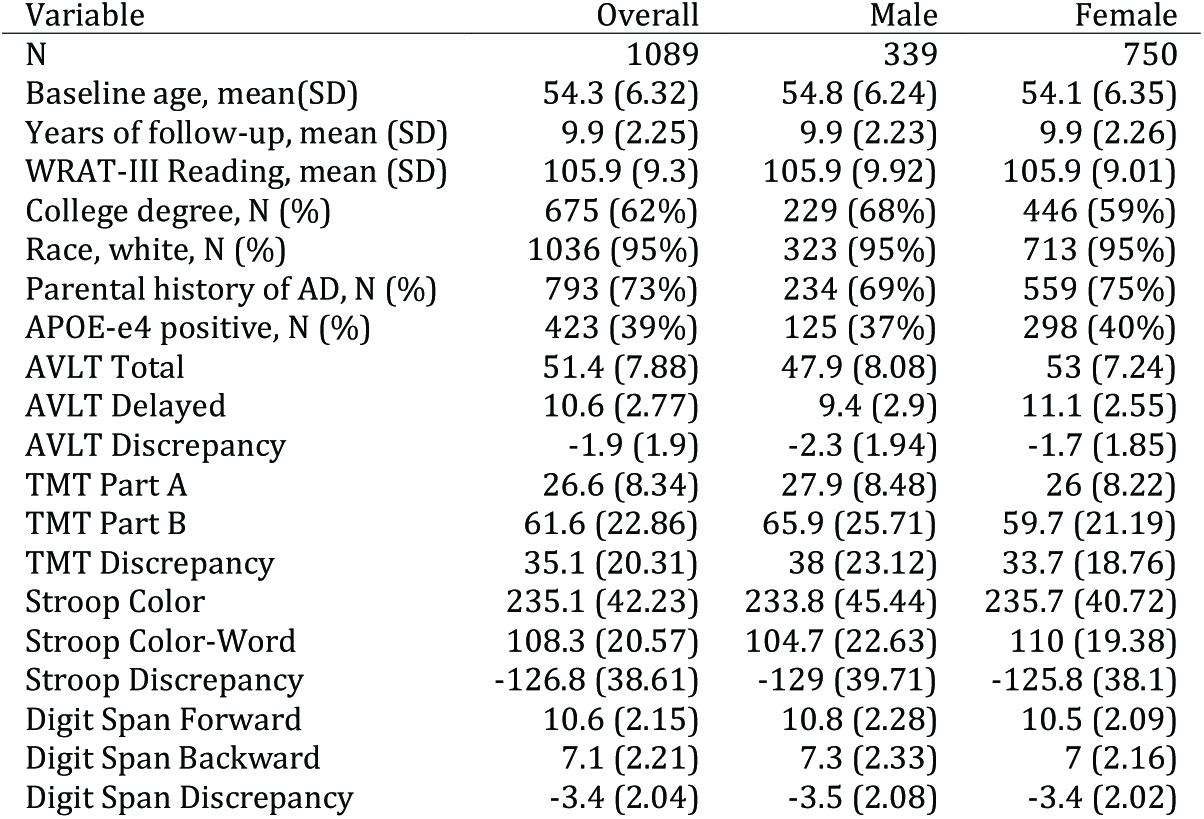
Baseline characteristics of sample, overall and separated by sex.

#### Unconditional standards

Coefficients for unconditional regression quantiles at median and 7th-percentile performance are listed in Table 2 for all outcomes. Notably, after adding demographic terms, the coefficients for age were near zero (indicating minimal change per year) in several models.

**Table 2.**
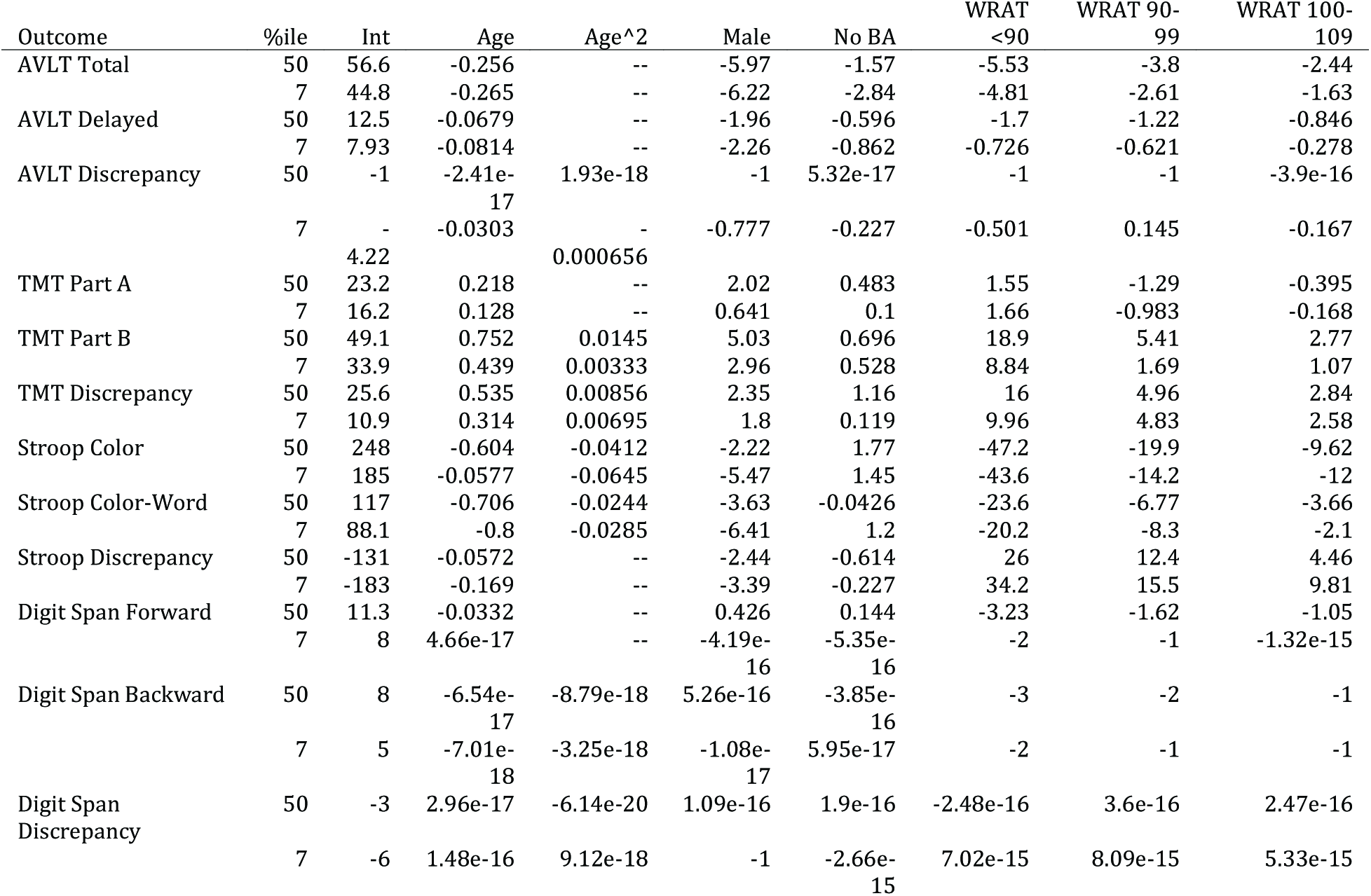
Regression coefficients for median and 7th-percentile unconditional models. Quadratic age terms were retained when nominally significant (p<.05) in preliminary modelsfor at least two quantiles. Models were otherwise unselected, in that all coefficients were retained regardless of significance.

#### Conditional standards

Coefficients for conditional regression quantiles at median and 7th-percentile performance are listed in Table 3. Among these models, the linear age coefficient was near zero for Digit Span Backward (median) and Stroop Discrepancy (7th percentile). Coefficients for practice indicated benefit from previous exposures, except for Stroop Color-Word.

**Table 3.**
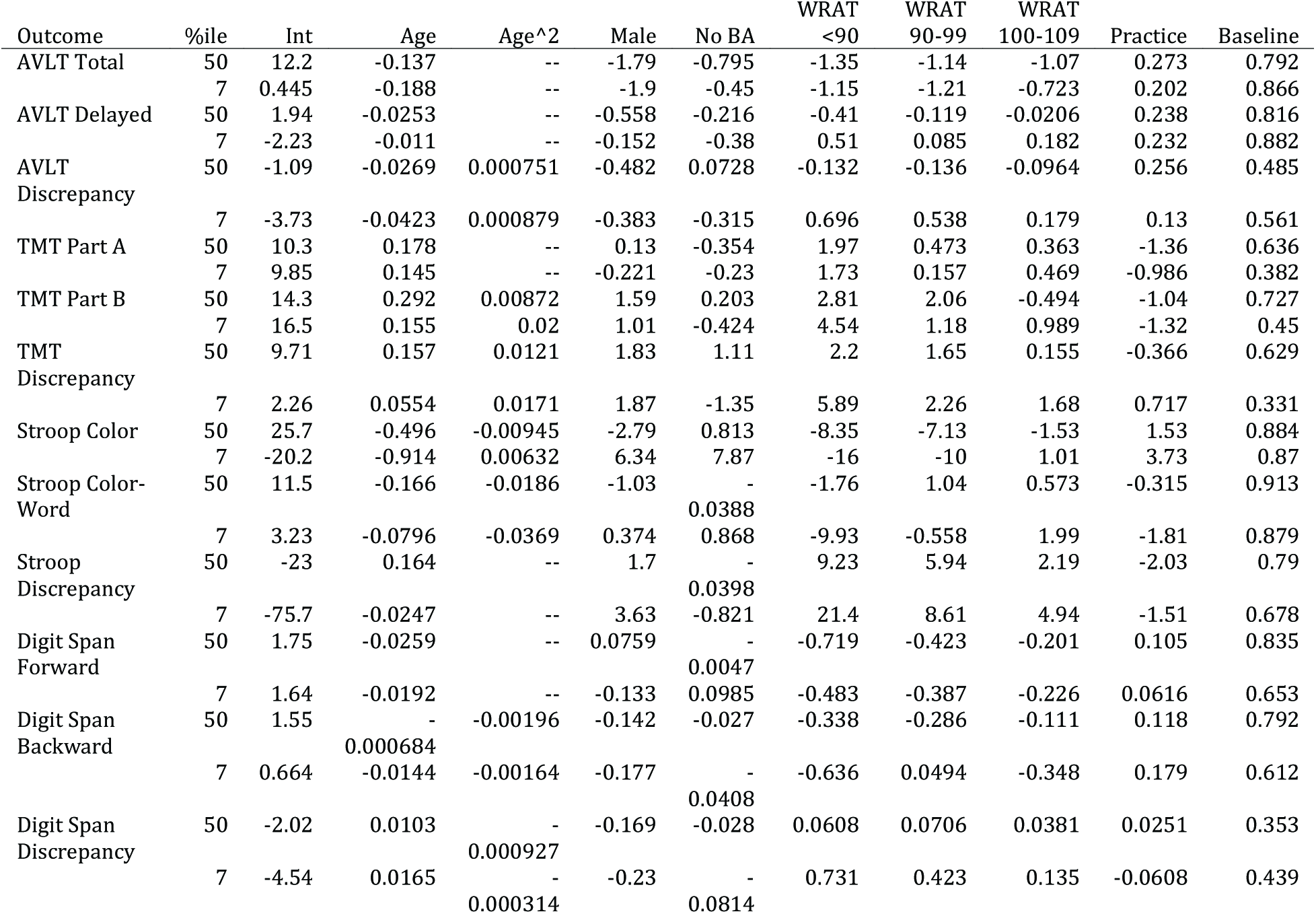
Regression coefficients for median and 7th-percentile conditional models.

#### Graphical display of example cases

The R programming platform was used to develop a graphical display of an individual participant’s performance superimposed on the unconditional standards with scores falling below our conditional 7th percentile cut-off being demarcated by a red circle. In Figure 1, longitudinal performance of three participants are shown for AVLT Total, AVLT Delayed, and Stroop Color-Word.

**Figure 1.**
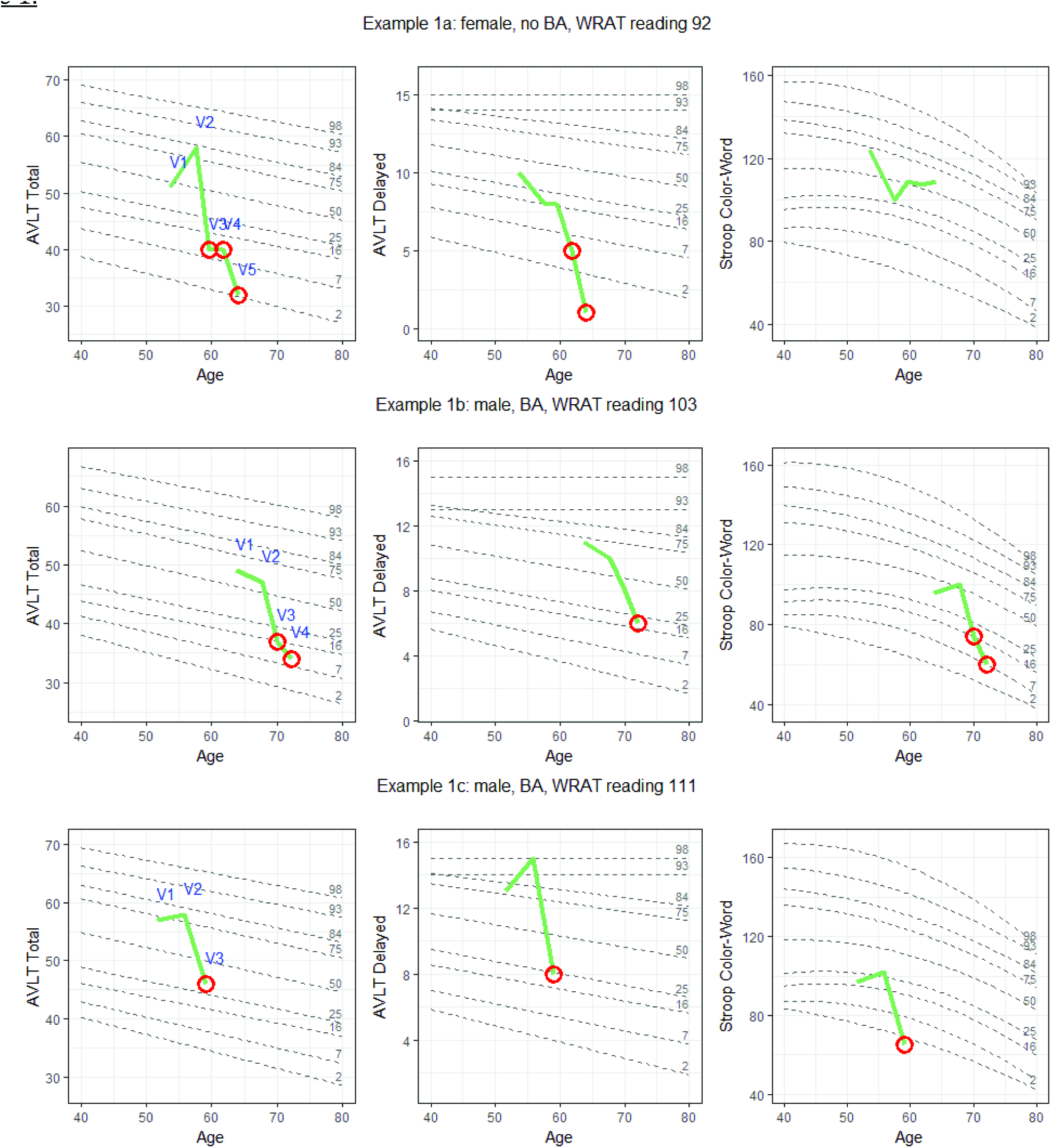
Longitudinal performance of three individuals (black) on each of three cognitive tests. Performances are plotted against demographically-adjusted unconditional standard lines for several percentiles (grey). Circles indicate abnormal conditional performance (ACP). Additional case details: Figure 1a: Full-scale IQ=107 (68th percentile); has been involved with caregiving for family members since enrollment (including a parent with AD from Visits 1 to 3); worked as a manager at enrollment and retired between Visits 3 and 4. No self-report of significant history of mental health problems; CES-D scores in normal range for all visits. Significant back pain at Visit 5 required a minor testing accommodation. Figure 1b: Full scale IQ 113 (88th percentile); retired teacher/coach at enrollment; history of depression and anxiety (began taking buproprion between Visits 3 and 4); most recent CES-D score was 34, indicating moderate depressive symptomatology; also reports hearing difficulty (evident in testing at Visit 4). Figure 1c: Full-scale IQ of 133 (99th percentile); works as a management consultant (fulltime at enrollment, part-time at and after Visit 2); no self-report of significant mental health history, and normal CES-D scores at all WRAP visits; stroke between Visits 2 and 3.

Figure 1a illustrates the performance of a woman with a parental history of AD (PH+) who enrolled in WRAP at age 53, had WRAT-3 reading standard score of 92, had some college education, and has been followed for five visits over ten years. She was judged to be cognitively normal via consensus conference for the first four visits, and given a diagnosis of MCI at Visit 5. However, she exhibited ACP on at least one test at Visit 3 (the first for which ACP information was available and more than four years previous to MCI diagnosis) and at all subsequent visits. The earliest test to show change was AVLT Total. By Visit 4, she was also exhibiting ACP on AVLT Delay, Trails Discrepancy, and Stroop Color. At Visit 5 (concurrent with her first clinical diagnosis), ACP emerged on AVLT Discrepancy.

Figure 1b illustrates the performance of a man with no parental history of AD (PH-) who enrolled in WRAP at age 63, had a WRAT-3 reading standard score of 103, had some graduate school training, and has been followed for four visits over eight years. He was judged at consensus conference to be cognitively normal for three visits and Early MCI at Visit 4 (i.e., subclinical deficits). However, he exhibited ACP on AVLT Total and Stroop Color-Word at Visit 3, two years previously, and on several tests at Visit 4, including AVLT Total, AVLT Delay, TMT Part A, TMT Part B, Stroop Color, and Stroop Color-Word.

Figure 1c illustrates the performance of another man (PH-) who enrolled in WRAP at age 51, had a WRAT-3 reading standard score of 111, had completed some graduate school and has been followed for three visits over seven years. His consensus conference diagnosis has been cognitively normal at all visits, but at Visit 3 (his most recent), he exhibited ACP on AVLT Total, AVLT Delay, TMT Part B, Stroop Color, and Stroop Color-Word.

### Validity of ACP

#### Concurrent cognitive status

Although the study is still ongoing with relatively few conversions to MCI and dementia, we can begin to assess validity by comparing ACP flags to cognitive statuses. Figure 2 depicts odds ratios and their confidence intervals from an ordinal GEE regression linking participants’ ACP flags (0=normal, 1=ACP) to their concurrent statuses (0=Cognitively Normal; 1=Early MCI; 2=MCI/Dementia). Odds ratio confidence intervals greater than 1 indicate increased risk of impairment among those with ACP vs normal conditional performance on that test. ACP on AVLT Total, AVLT Delayed, AVLT Discrepancy, TMT Part A, TMT Part B, Stroop Color, and Stroop Color-Word were each associated with higher odds of a concurrent cognitive status indicating impairment (Early MCI or MCI/Dementia). No relationships were observed with any flag based on Digit Span. Confidence intervals for most discrepancy scores overlapped zero, and no discrepancy score offered more information than its component scores.

**Figure 2.**
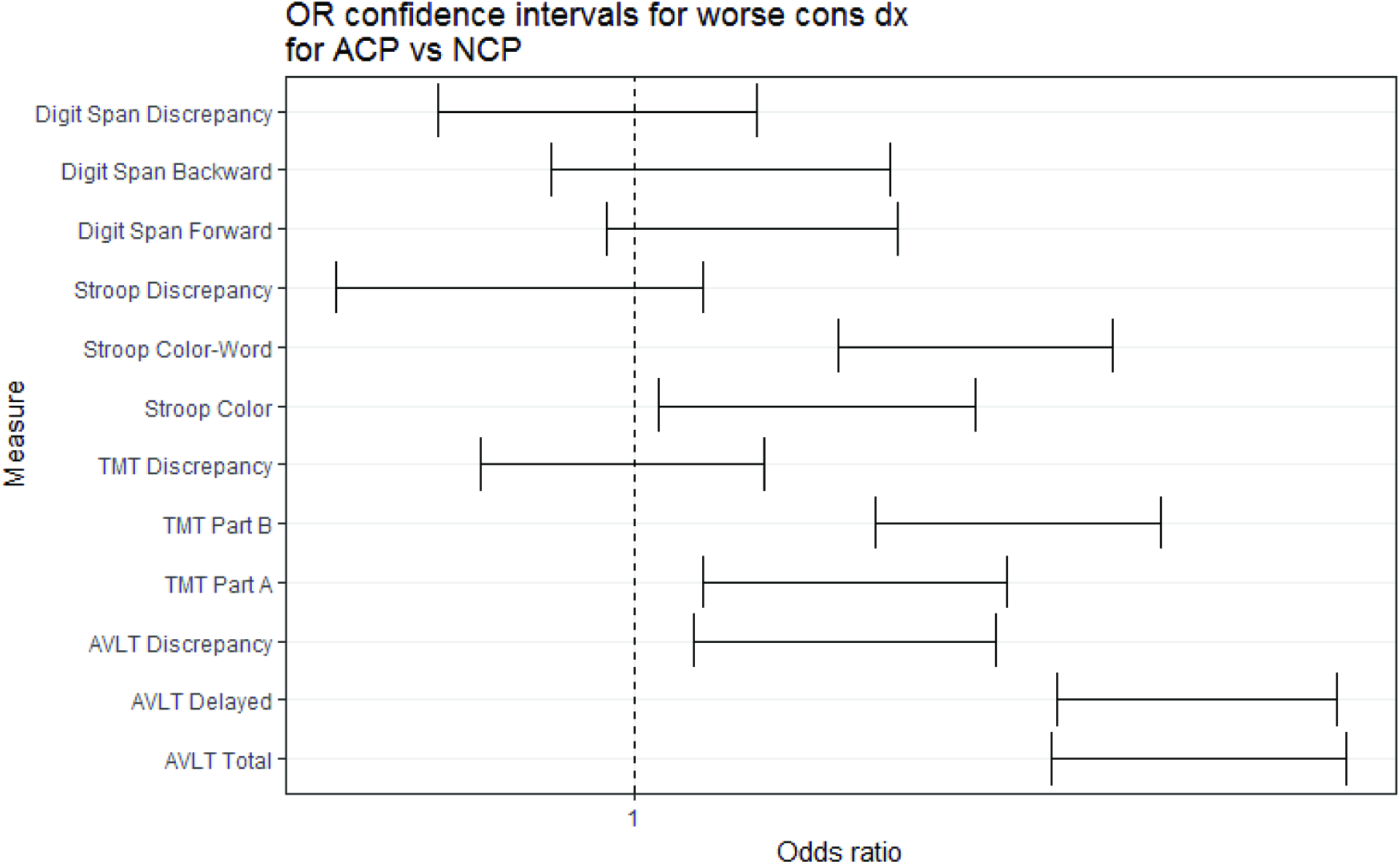
Confidence intervals on odds ratio estimates for ordinal regression predicting cognitive status from concurrent ACP.

#### Prospective cognitive status

We can also begin to ask whether ACP on a given test at an early visit improves prediction of later cognitive outcomes, after controlling for AUP on the same test. Among the subset that was cognitively normal at Visit 3 and had a final cognitive status at Visit 4 or later (N=709), we examined whether first available ACP (at Visit 3) was predictive of last available cognitive status (at visit 4, 5, or 6; 0=Cog Normal (N=651), 1=EarlyMCI (N=53), 1=MCI/Dementia (N=5)), controlling for Visit 3 AUP. In no analysis was this predictive relationship significant. AUP itself was predictive of final diagnosis for AVLT Delayed and TMT part B. Results are summarized in Table 4.

**Table 4.** Summary of ordinal regression results predicting most recent cognitive status from Visit 3 ACP and AUP. Reported p-values have been adjusted using the Benjamini-Hochberg procedure.

| Outcome | ACP odds ratio (CI) | ACP p-value | AUP odds ratio (CI) | AUP p-value |
| --- | --- | --- | --- | --- |
| AVLT Total | 1.21 (0.36-4.07) | 0.818 | 5.24 (1.43-19.26) | 0.056 |
| AVLT Delayed | 2.02 (0.69-5.94) | 0.818 | 5.8 (1.85-18.14) | 0.009 |
| AVLT Discrepancy | 1.47 (0.41-5.35) | 0.818 | 1.93 (0.53-7.05) | 0.218 |
| TMT Part A | 0.67 (0.13-3.4) | 0.818 | 1.2 (0.23-6.23) | 0.979 |
| TMT Part B | 1.22 (0.38-3.91) | 0.818 | 5.24 (1.56-17.62) | 0.014 |
| TMT Discrepancy | 1.97 (0.51-7.64) | 0.818 | 0.35 (0.08-1.6) | 0.402 |
| Stroop Color | 1.18 (0.36-3.84) | 0.818 | 1.18 (0.36-3.84) | 0.740 |
| Stroop Color-Word | 1.54 (0.53-4.52) | 0.818 | 1.87 (0.58-6.08) | 0.288 |
| Stroop Discrepancy | 1.84 (0.65-5.15) | 0.818 | 1 (0.29-3.41) | 0.693 |
| Digit Span Forward | 0.87 (0.27-2.82) | 0.818 | 1.82 (0.76-4.35) | 0.388 |
| Digit Span Backward | 0.55 (0.14-2.11) | 0.818 | 1.65 (0.61-4.47) | 0.693 |
| Digit Span Discrepancy | 0.62 (0.1-3.82) | 0.818 | 0.62 (0.1-3.82) | 0.400 |

In a secondary analysis on the larger subset whose cognitive status was not clinically impaired (inclusive of normal and early MCI at visit 3; N=830), we examined whether first available ACP and concurrent AUP on a given test were significant predictors of the last available clinical status (0=Cog Normal or EarlyMCI (N=815), 1=MCI/Dementia (N=15)). As in the ordinal results, clinical status was not related to Visit 3 ACP for any of the neuropsychological measures, and only related to Visit 3 AUP for AVLT Delayed.

### Subjective memory complaints and informant reports

After adjustment for multiple comparisons, Likert-scale self-ratings of memory showed significant relationships with ACP on AVLT Total and AVLT Delayed, and with AUP on AVLT Delayed and TMT part B (Supplemental Table 1; Figure 3). Similarly, participants’ responses to the binary question, “Do you think you have a problem with your memory?” were significantly related to ACP on AVLT Total and AVLT Delayed, but not to ACP in other non-memory domains (Supplemental Table 2). Finally, models using informant scores on the IQCODE to predict concurrent ACP are summarized in Table 5. Whereas subjective memory related most strongly to ACP on memory measures, IQCODE related instead to ACP on the challenge-conditions from three different tests: AVLT Delayed, TMT part B, and Stroop Color-Word.

**Figure 3.**
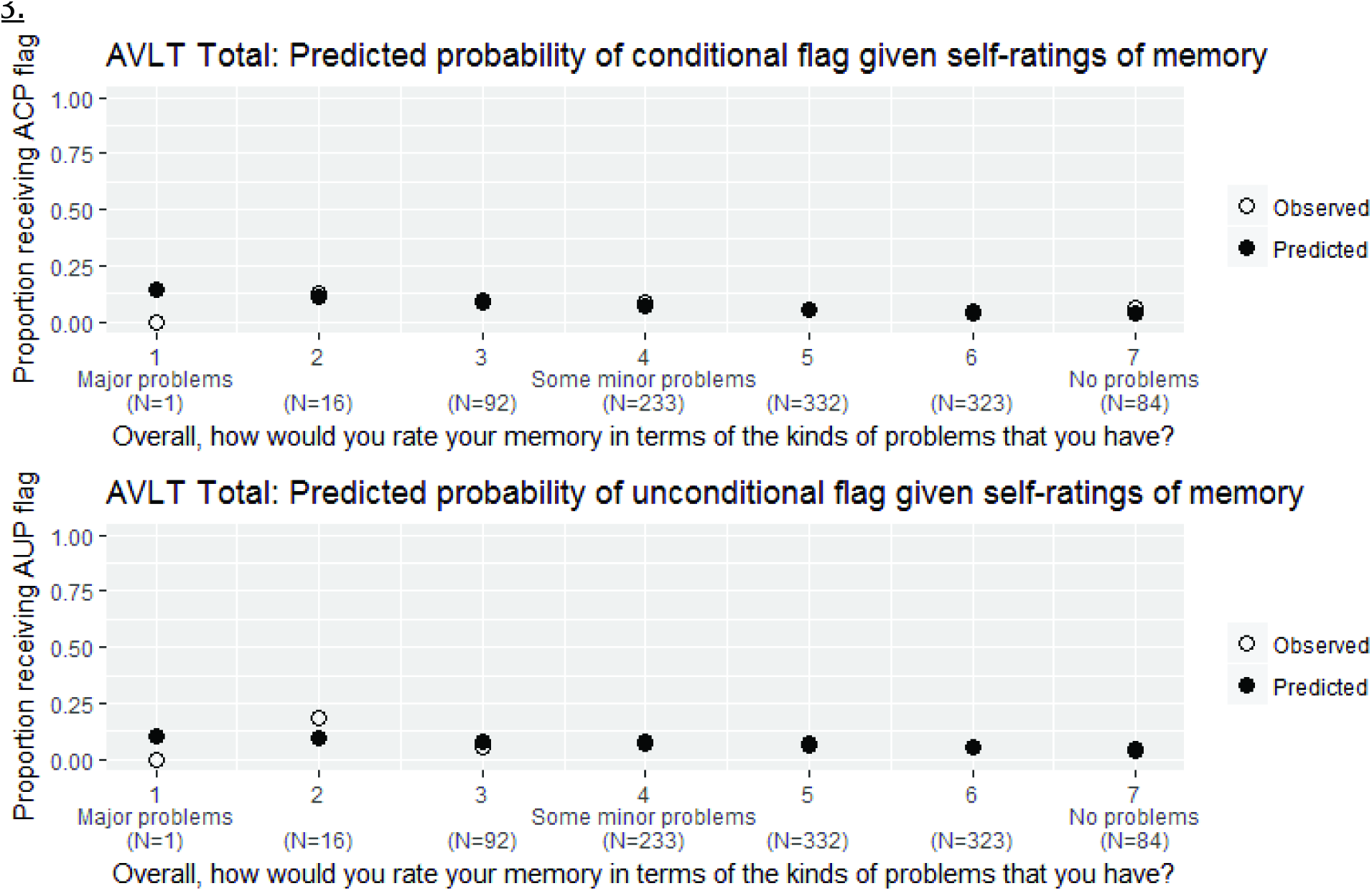
Predicted probability plots for models of ACP (first panel) and AUP (second panel) on AVLT Total as a function of subjective memory performance (x-axis). The linear predictor is shown in black dots, and the observed proportions at Visit 3 in open circles; the label indicates the total N observed at Visit 3 for each x-value.

**Table 5.** Summary of GEE model predicting ACP from informant reports of cognitive functioning (IQCODE). Higher scores on this measure indicate worse functioning; accordingly, odds ratios represent change in risk associated with one-point increase in reports of worsened function. Reported p-values have been adjusted using the Benjamini-Hochberg procedure.

| Outcome | ACP odds ratio (CI) | ACP LR Chi-squared | ACP p-value |
| --- | --- | --- | --- |
| AVLT Total | 1.02 (0.96-1.08) | 0.29 | 0.683 |
| AVLT Delayed | 1.19 (1.07-1.32) | 10.15 | 0.009 |
| AVLT Discrepancy | 1.06 (1-1.14) | 3.61 | 0.172 |
| TMT Part A | 1.03 (0.96-1.09) | 0.61 | 0.683 |
| TMT Part B | 1.12 (1.03-1.22) | 7.37 | 0.027 |
| TMT Discrepancy | 0.99 (0.95-1.03) | 0.35 | 0.683 |
| Stroop Color | 1 (0.96-1.04) | 0.01 | 0.904 |
| Stroop Color-Word | 1.15 (1.06-1.24) | 12.44 | 0.005 |
| Stroop Discrepancy | 1.03 (0.98-1.07) | 1.24 | 0.638 |
| Digit Span Forward | 1.01 (0.97-1.05) | 0.24 | 0.683 |
| Digit Span Backward | 1.02 (0.96-1.07) | 0.30 | 0.683 |
| Digit Span Discrepancy | 1.02 (0.97-1.08) | 0.54 | 0.683 |

### Summary statistics: numbers flagged under each method

Table 6 contains numbers of participants identified at Visit 3 as exhibiting ACP, AUP, or both. If regression quantiles are well-constructed, approximately 6-7% of measurements should be flagged as either ACP or ACP+AUP; similarly, 6-7% of measurements should be flagged as either AUP or ACP+AUP. Conditional models performed as expected, flagging 6-7% of observations at Visit 3 for all neuropsychological outcomes. Unconditional models were a bit more variable, generally flagging 5-8% of observations, but more for Digit Span Forward (10%) and Digit Span Backward (10.1%). We see substantial overlap between the categories, with disproportionate numbers receiving both flags; however, for most of the tests, AUP and ACP each detect evidence of poor performance when the other does not.

**Table 6.** Frequency of ACP and AUP flags at Visit 3, by subtest.

| Outcome | Normal | AUP Only | ACP Only | ACP and AUP |
| --- | --- | --- | --- | --- |
| AVLT Total | 977 (89.8%) | 44 (4.0%) | 39 (3.6%) | 28 (2.6%) |
| AVLT Delayed | 972 (89.3%) | 48 (4.4%) | 32 (2.9%) | 36 (3.3%) |
| AVLT Discrepancy | 998 (91.7%) | 19 (1.7%) | 20 (1.8%) | 51 (4.7%) |
| TMT Part A | 992 (91.6%) | 24 (2.2%) | 33 (3.0%) | 34 (3.1%) |
| TMT Part B | 976 (90.2%) | 34 (3.1%) | 42 (3.9%) | 30 (2.8%) |
| TMT Discrepancy | 976 (90.5%) | 32 (3.0%) | 22 (2.0%) | 48 (4.5%) |
| Stroop Color | 962 (89.3%) | 45 (4.2%) | 35 (3.2%) | 35 (3.2%) |
| Stroop Color-Word | 963 (89.8%) | 38 (3.5%) | 40 (3.7%) | 31 (2.9%) |
| Stroop Discrepancy | 970 (90.6%) | 29 (2.7%) | 40 (3.7%) | 32 (3.0%) |
| Digit Span Forward | 950 (87.2%) | 66 (6.1%) | 31 (2.8%) | 42 (3.9%) |
| Digit Span Backward | 951 (87.3%) | 68 (6.2%) | 27 (2.5%) | 43 (3.9%) |
| Digit Span Discrepancy | 998 (91.6%) | 23 (2.1%) | 17 (1.6%) | 51 (4.7%) |

## Discussion

This paper presents a novel method for detection of preclinical cognitive changes based on a method first developed for anthropometric indices, namely, longitudinal reference curves (Bowditch, 1891; Healy, 1974). Unconditional and conditional standards for an age range of 40-75 were derived from longitudinal assessments of the WRAP cohort of cognitively healthy individuals enriched for parental history of AD dementia. Both sets of standards incorporate sex, literacy, age, and education in development of equations representing percentiles spanning from low to high functioning. We illustrate the utility of these standards by plotting longitudinal performance of three WRAP participants superimposed on the unconditional percentiles, with highlighted scores representing potentially troubling within-person change according to conditional standards. In these cases, abnormal conditional performance on one or more tests heralded an abnormal consensus conference diagnosis associated with MCI or probable AD (cases 1 and 2) or stroke (case 3). We found that test performance outside of expected ranges based on unconditional and conditional standards for cognitive change were associated with subjective and informant reports of cognitive decline, and a concurrent consensus conference diagnosis of cognitive impairment.

The application of quantile regression methods to these tests represents a significant extension of earlier work presenting unconditional and conditional standards for MMSE (Cheung et al., 2015). Compared to WRAP, their sample was older (mean baseline age=65) and had less follow-up data (maximum of three visits). By extending the method to tests sensitive to early cognitive change, we can begin to evaluate its utility in populations most likely to benefit from prevention efforts.

The development of conditional standards directly addresses new criteria for identifying the earliest stages of decline. In the model of the Alzheimer’s pathophysiological continuum currently in development by Jack and colleagues (2017), Stage 2 corresponds to a drop in cognitive performance that has not yet crossed objective thresholds for cognitive impairment. Current clinical practice does not have an agreed-upon method for identifying worrisome change. We propose the method described in this paper as a starting point to operationalize such decline. Follow-up analyses will be done using different operationalizations for decline (e.g., multiple tests with ACP or different centile threshold for “low”) when more of the sample has converted to dementia, and when biomarkers are available on enough of the sample to evaluate the utility of conditional standardsin the context of Jack and colleagues’ (2017) ATN framework.

Our method is similar in some ways to earlier regression-based methods of identifying reliable change (Crawford et al., 2012; Maassen et al., 2009). The major difference in the quantile regression approach is that it more naturally handles tests whose distributions cannot be expected to be normal, such as the MMSE (e.g. as in Cheung et al., 2015), and situations in which relationships between predictors and outcomes varies by quantile, as may well be the case for age trajectories on cognitive tests. Future work will explore the relationship between these two methods on the same dataset to better understand the degree of overlap between them.

Our data suggest that the conditional standards may be sensitive to subjective changes in memory, a construct whose utility has been debated (Roberts et al., 2009). The existence of a modest relationship suggests that our approach has face validity, as both measures attempt to estimate cognitive change from some presumably healthy baseline. It is possible that our conditionally-poor performers represent a mix of people who are aware of their deficit, and people whose decline is already advanced enough that they are experiencing anosognosia (Roberts et al., 2009; Vannini et al., 2017). Future analyses will examine these possibilities.

We included discrepancy scores in our data because they have been thought by some clinicians to be especially sensitive measures (Hayden et al., 2014; Jacobson, Delis, Bondi, & Salmon, 2002; though see also Smith, Ivnik, & Lucas, 2008). In our dataset, however, knowledge of widening gaps between the selected test pairs did not improve on what was learned from component scores (Figure 2). The reliability of discrepancy scores is bounded by the reliability of the two component scores and their correlation (Crawford, Sutherland, & Garthwaite, 2008), which limits their longitudinal usefulness.

### Limitations and future directions

While promising, this work has limitations. These standards reflect the performance of a single, non-population-based sample of highly educated, mostly white individuals. The 7th percentile threshold we specified for abnormality, while consistent with past conventional approaches, may not be the most useful for all contexts. In addition, there may be information in either the relationship between concurrent ACP flags on different domains, or in the temporal sequencing of flags within a domain, which we have not explored in these data. Our technique shares with regression-based norms of the past the need for a large longitudinal dataset on which standards for a given test can be developed. Our use of a multiple-visit baseline limits the immediate clinical application of this technique, since ACP as we have defined it cannot be determined before a patient’s third clinic visit. The benefit of our approach is that it reduces spurious identification of patients whose apparent change represents regression toward the mean. Finally, we caution that the conditional and unconditional standards presented here only capture test performance and thus should not be used in isolation. As always, clinical judgement is central to making any clinical or research diagnoses based on the totality of information available.

These limitations may be addressed in future analyses in one or more of the following ways. Data from multiple sources can be pooled to increase representativeness of standards and generate training and validation sets. With such a dataset, it could also be useful to compare robust longitudinal standards, obtained by removing individuals with extremely unusual longitudinal performance from the standards development sample, to the conventional ones described in this paper (cf. Clark et al., 2016; De Santi et al., 2008; Koscik et al., 2014). In future work, we will examine the usefulness and stability of “provisional” conditional standards for the second visit. Future analyses will also explore additional operationalizations of preclinical decline using conditional and unconditional centiles (e.g., using different centile thresholds for defining significant change or low performance on a given test, using tallies of numbers of tests showing low performance or abnormal changes, etc.), to determine how best to leverage performance relative to the standards on multiple tests to create robust indicators of cognitive change or dementia risk.

### Conclusions

In this paper, we have presented application of quantile regression to development of unconditional and conditional (longitudinal) standards for performance on a variety of cognitive measures that are sensitive to cognitive decline associated with AD dementia. The graphical interface offers a clinically intuitive visualization of an individual’s cognitive performance across measures and over time and clearly identifies tests demonstrating potentially troubling within person change from baseline. Such methods are needed in order to operationalize preclinical declines in AD and other dementias.

## Acknowledgements

This research was supported by the National Institutes of Health awards (R01 AG027161, R01 AG021155, R01 AG054059, P50 AG033514, and UL1 TR000427, and by donor funds including the Wisconsin Alzheimer’s Institute Lou Holland Fund and contributions from anonymous donors. Portions of this research were supported by resources at the Wisconsin Alzheimer’s Institute, the Wisconsin Alzheimer’s Disease Research Center, and the Geriatric Research Education and Clinical Center of the William S. Middleton Memorial Veterans Hospital, Madison, WI. Any opinions, findings, and conclusions or recommendations expressed in this material are those of the authors(s) and do not necessarily reflect the views of the NIH or the Veterans Administration. The authors gratefully acknowledge the WRAP study team who have carefully acquired the longitudinal data and the WRAP participants who make this research possible.. The authors of this manuscript have no conflicts of interest to report.

